# Plasma amyloid-β homeostasis is associated with Body Mass Index and weight loss in people with overweight and obesity

**DOI:** 10.1101/2022.05.31.494083

**Authors:** Emily S. Brook, Zachary J. D’Alonzo, Virginie Lam, Dick Chan, Satvinder Singh Dhaliwal, Gerald F. Watts, John C. L Mamo, Ryusuke Takechi

## Abstract

**BACKGROUND:** Obesity is linked to a higher incidence of Alzheimer’s disease (AD). Studies show that plasma amyloid-β (Aβ) dyshomeostasis, particularly low 42/40 ratio indicates a heightened risk for developing AD. However, the relationship between body mass index (BMI) and circulating plasma Aβ has not been extensively studied.

**OBJECTIVE:** We hypothesised that people with a high BMI have altered plasma Aβ homeostasis compared with people with a lower BMI. We also tested whether reducing BMI by calorie-restriction could normalise plasma concentrations of Aβ.

**METHODS:** Plasma concentrations of Aβ_40_, Aβ_42_ and Aβ_42/40_ ratio were measured in 106 participants with BMIs classified as lean, overweight, or obese. From this cohort, twelve participants with overweight or obese BMIs entered a 12-week calorie-restriction weight loss program. We then tested whether decreasing BMI affected plasma Aβ concentrations.

**RESULTS:** Plasma Aβ_42/40_ ratio was 17.54% lower in participants with an obese BMI compared to lean participants (p<0.0001), and 11.76% lower compared to participants with an overweight BMI (p<0.0001). The weight loss regimen decreased BMI by an average of 4.02% (p=0.0005) and was associated with a 6.5% decrease in plasma Aβ_40_ (p=0.0425). However, weight loss showed negligible correlations with plasma Aβ_40_, Aβ_42_ and Aβ_42/40_ ratio.

**CONCLUSION:** Obesity is associated with aberrant plasma Aβ homeostasis which may be associated with an increased risk for AD. Weight loss appears to lower Aβ_40_, but large-scale longitudinal studies in addition to molecular studies are required to elucidate the underlying mechanisms of how obesity and weight loss influence plasma Aβ homeostasis.

## INTRODUCTION

Alzheimer’s disease (AD), the most common form of dementia, is characterised by progressive cognitive decline and the abnormal deposition of amyloid-β (Aβ) in the brain parenchyma [1]. A significant risk factor for AD is obesity, which is defined as body mass index (BMI) being ≥30 kg/m^2^ [2, 3]. Obesity in people aged between 40-45 years has been associated with a three-fold higher risk of developing AD in later-life [4]. Additionally, obesity was reported to significantly accelerate the onset of AD and double the likelihood of Aβ deposition in the brain [5, 6]. More recently, overweight and obese BMIs were found to be associated with detrimental changes to brain structure, such as increased white matter oedema and low hippocampal volume [7]. These data collectively suggest that an increased BMI may have a severe impact on cerebral Aβ overload, AD onset and progression, in addition to overall brain health. However, the exact underlying mechanisms linking obesity with AD remain unknown.

Many studies have evaluated plasma levels of Aβ_40_ and Aβ_42_ as predictors of AD or cognitive decline [8]. Aβ_40_ and Aβ_42_ are generated from the sequential proteolytic cleavage of Aβ precursor protein (AβPP) by β-site AβPP cleaving enzyme-1 (BACE1) and γ-Secretase [9]. Aβ_40_ is the most physiologically abundant species of amyloid, whilst Aβ_42_ is regarded as primarily pathogenic as it is prone to aggregation and eventually forms insoluble fibrillary structures which are the core component of amyloid plaques [10, 11]. Recent findings show that the dyshomeostasis of plasma Aβ, particularly low plasma Aβ_42/40_ ratio can predict the risk of AD at an accuracy of ~90% [12–19]. However, few studies to date have investigated the association between BMI, weight loss and plasma Aβ homeostasis [20, 21], although obesity is a significant modifiable risk factor for AD.

In this pilot study, we investigated whether BMI shows any significant associations with plasma Aβ homeostasis. Furthermore, in individuals with BMIs classified as overweight or obese, we tested whether reducing BMI through a calorie restriction weight loss regime could modulate plasma Aβ homeostasis. The outcomes of this study may provide new insights into the mechanisms whereby obesity increases brain amyloidosis and risk of AD and may offer potential preventative opportunity with weight loss.

## MATERIALS AND METHODS

### PARTICIPANTS

To investigate the association between BMI and plasma Aβ homeostasis, we recruited 106 people who either had a normal, overweight, or an obese BMI. BMI cut-offs were defined according to the World Health Organisation: lean weight as 18.50-24.99 kg/m^2^, overweight as 25.00-29.99 kg/m^2^, and obese as ≥30.00 kg/m^2^ [22]. No participants had an underweight BMI (≤18.50 kg/m^2^). All participants were Caucasian, and those with familial hypercholesterolemia, a history of cardiovascular disease or used antidiabetic medication were excluded from the study. In addition, participants with APOE/E2 genotype were excluded. Aside from ageing, APOE genotype is the most significant genetic risk factor for AD, with APOE2/E2 conferring for reduced AD risk as detailed in [23]. This study was approved by the Ethics Committee of Royal Perth Hospital (EC 2010/074), a national ethics committee (Bellberry Ltd, Eastwood, South Australia, 2014-03-151) and Curtin University (HRE2021-0192). Informed consent was obtained from each participant.

### SAMPLE COLLECTION AND STUDY DESIGN

After a 14-hour fast, baseline whole venous blood was collected in EDTA tubes from all 106 participants and immediately centrifuged at 1500 x *g* for 15 min at 4°C to obtain plasma. Plasma was collected and stored at −80°C for future evaluation. At this visit, we also recorded all participants’ weight and height for their BMI calculation.

### WEIGHT LOSS PROGRAM

Following the plasma sample and BMI data collection, all participants who had an overweight or obese BMI were invited to participate in a 12-week calorie restriction weight loss program as reported previously [23], and 12 chose to participate. During this weight loss regime, the participants’ caloric intake was restricted to achieve a deficit of ~1440 kJ per day compared to their diets before the commencement of weight loss. Dietary intake was evaluated for energy and major nutrients by FoodWorks 2007 (Xyris Software, Brisbane, Australia). All participants were reviewed fortnightly and were requested to maintain their usual level of physical activity. All dietary assessments and recommendations were conducted by a registered dietician. After completing the weight loss program, fasting plasma was collected, and BMI was recorded for the 12 participants.

### QUANTIFICATION OF PLASMA AB_40_ AND AB_42_

Plasma concentrations of Aβ_40_ and Aβ_42_ were measured at baseline (*n*=106) and following the 12-week weight loss program (*n*=12) using ELISA (Wako Chemical, Tokyo, Japan), according to the manufacturer’s instruction. Briefly, samples were assays in singlets and prepared in a 1 in 4 dilution with the kit provided diluent. All samples and kit-provided standards were loaded onto the antibody coated microplate and incubated overnight at 4°C. Following incubation, HRP-conjugated monoclonal antibody was added to the appropriate wells, and plates were incubated in the dark. Plates detecting Aβ_40_ were incubated for 2 hours, whilst plates detecting Aβ_42_ were incubated for 1 hour. Subsequently, 100 μL TMB was added to each well. The plates were incubated for a further 30 minutes in the dark, and STOP solution was added before the absorbance was measured at 450 nm using EnSpire™ Multimode Plate Reader (PerkinElmer, Waltham, MA, USA).

### STATISTICAL ANALYSIS

Statistical analyses were performed using GraphPad Prism 9.3.1 for Windows (GraphPad Software, CA, USA) and IBM SPSS Statistics for Windows, Version 27.0. (IBM Corp, NY, USA). The distribution of all variables was assessed by D’Agostino and Shapiro-wilk normality test. Unpaired t tests were used to compare variables of the three BMI groups. Where data was not normally distributed, intergroup comparisons were assessed using the Mann-Whitney test. For the calorie restriction part of the study, paired t tests were used to compare the difference from baseline to week-12. Where data was not normally distributed, changes were analysed using the Wilcoxon test. For correlation analyses, we used Pearson or Spearman correlation coefficient tests depending on the data normality, as well as principal component analysis (PCA). Where indicated, multivariable regression model was used to adjust data for age, age squared and sex. Significance was defined as *p*<0.05.

## RESULTS

The characteristics of the cohort are summarised in Table 1. Overall, participants were aged between 18 to 66 years. Individuals with an obese BMI were significantly older compared to participants with a lean or overweight BMI, whilst the average participant age of the lean and overweight groups were similar. Majority of the participants were male, accounting for 92.67% of the overall cohort.

**Table 1:** Cohort baseline characteristics. All variables are presented as mean ± SD with range. Non-matching superscript lettering indicates significant difference. Alphabetical letters indicate statistical significance at *p*< 0.05

| **BMI Groups** | **Lean**<br>(n=42) | **Overweight**<br>(n=41) | **Obese**<br>(n=23) |
| --- | --- | --- | --- |
| Age (years) | 32.40 ± 11.24<sup>a</sup> | 35.88 ± 12.66<sup>a</sup> | 57.57 ± 9.80<sup>b</sup> |
| (range) | (18-54) | (19-44) | (29-66) |
| Gender (% Male) | 100 | 95.12 | 60.87 |
| BMI (kg/m<sup>2</sup>) | 22.91 ± 1.40<sup>c</sup> | 27.25 ± 1.58<sup>d</sup> | 33.96 ± 5.12<sup>e</sup> |
| (range) | (19.29-24.99) | (25.00-29.88) | (30.04-51.07) |

The average plasma concentration of Aβ_40_ was significantly higher in people with an obese BMI compared to those with lean or overweight BMI by 41.7% and 36.89% respectively, whilst there was no significant difference observed between the lean and overweight groups (Fig 1A). Moreover, plasma abundance of Aβ_42_ was 12.51% higher in the obese group compared to overweight group (*p*= 0.0404); however, there were no other significant intergroup comparisons (Fig 1B). Lastly, plasma Aβ_42/40_ ratio showed a significant sequential decline as BMI categories progressed from lean, to overweight, to obese. Specifically, plasma Aβ_42/40_ ratio was 32.65% and 25.74% lower in the obese group compared to the lean and overweight BMI groups respectively, and 7.05% lower in the overweight cohort compared to the lean group (Fig 1C).

**Fig. 1.**
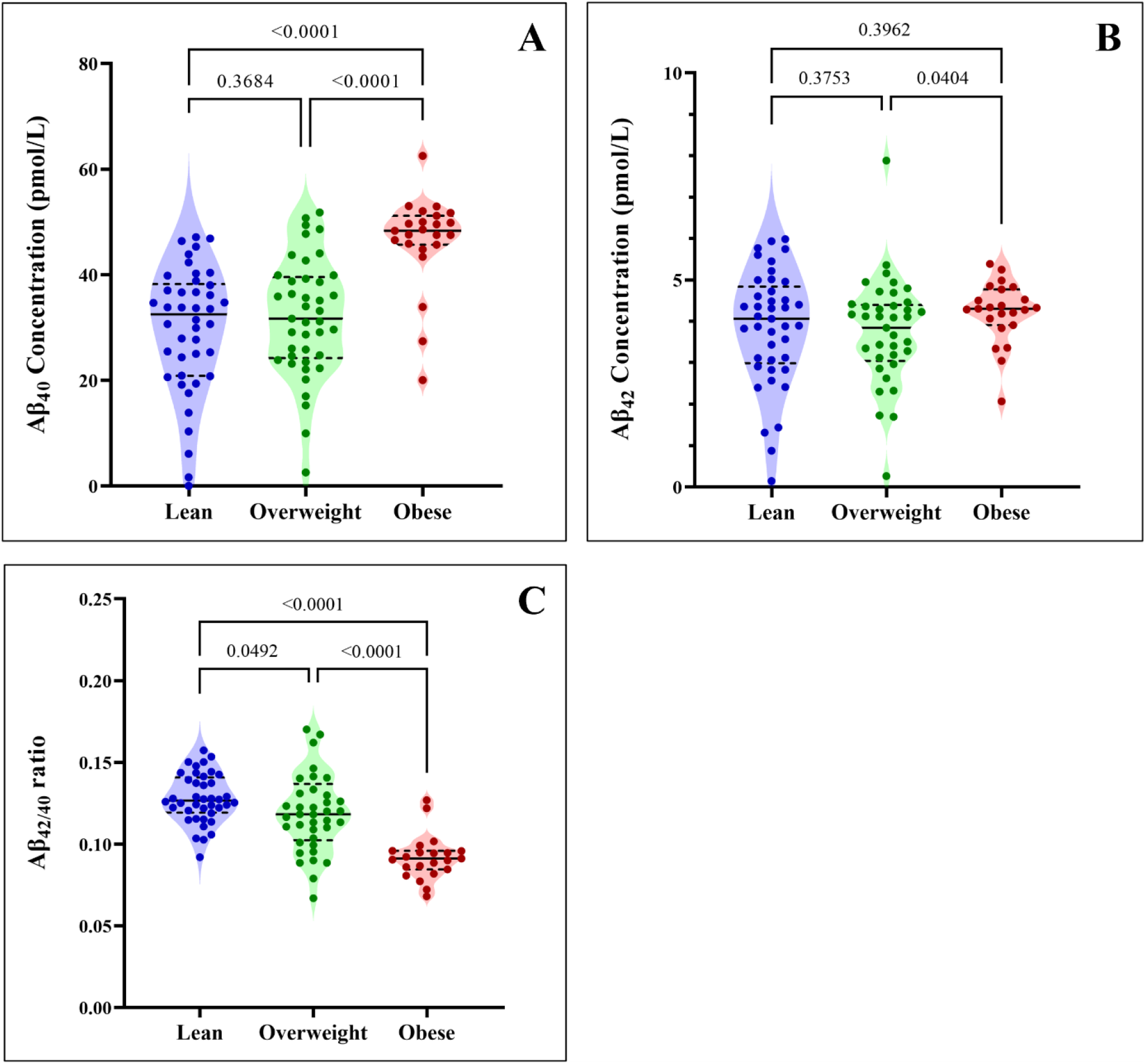
Violin plots show the plasma concentrations of A) Aβ_40_, B) Aβ_42_, and C) Aβ_42/40_ in people with lean, overweight, or obese BMIs. The median value is indicated by the solid line, and the quartiles are represented by the dashed lines.

To further evaluate the relationship between plasma Aβ and BMI, we performed simple linear correlation analysis (Fig 2). BMI showed a significant positive correlation with plasma Aβ_40_ concentration (Fig 2A), while no significant association was observed between BMI and Aβ_42_ (Fig 2B). Additionally, we identified a moderate negative correlation between BMI and Aβ_42/40_ ratio (Fig 2C). Consistently, the PCA loading plot showed a strong negative relationship between BMI and Aβ_42/40_ as indicated by the large angle and opposing direction of the two variables. However, Aβ_40_ and Aβ_42_ *per se* appeared as independent factors as indicated by their near orthogonal directions in relation to BMI (Fig 2D). In addition, the PCA scores plot displayed evident clustering of lean, overweight, and obese populations, with obesity showing negative correlation with Aβ_42/40_ ratio (Fig 2E).

**Fig. 2.**
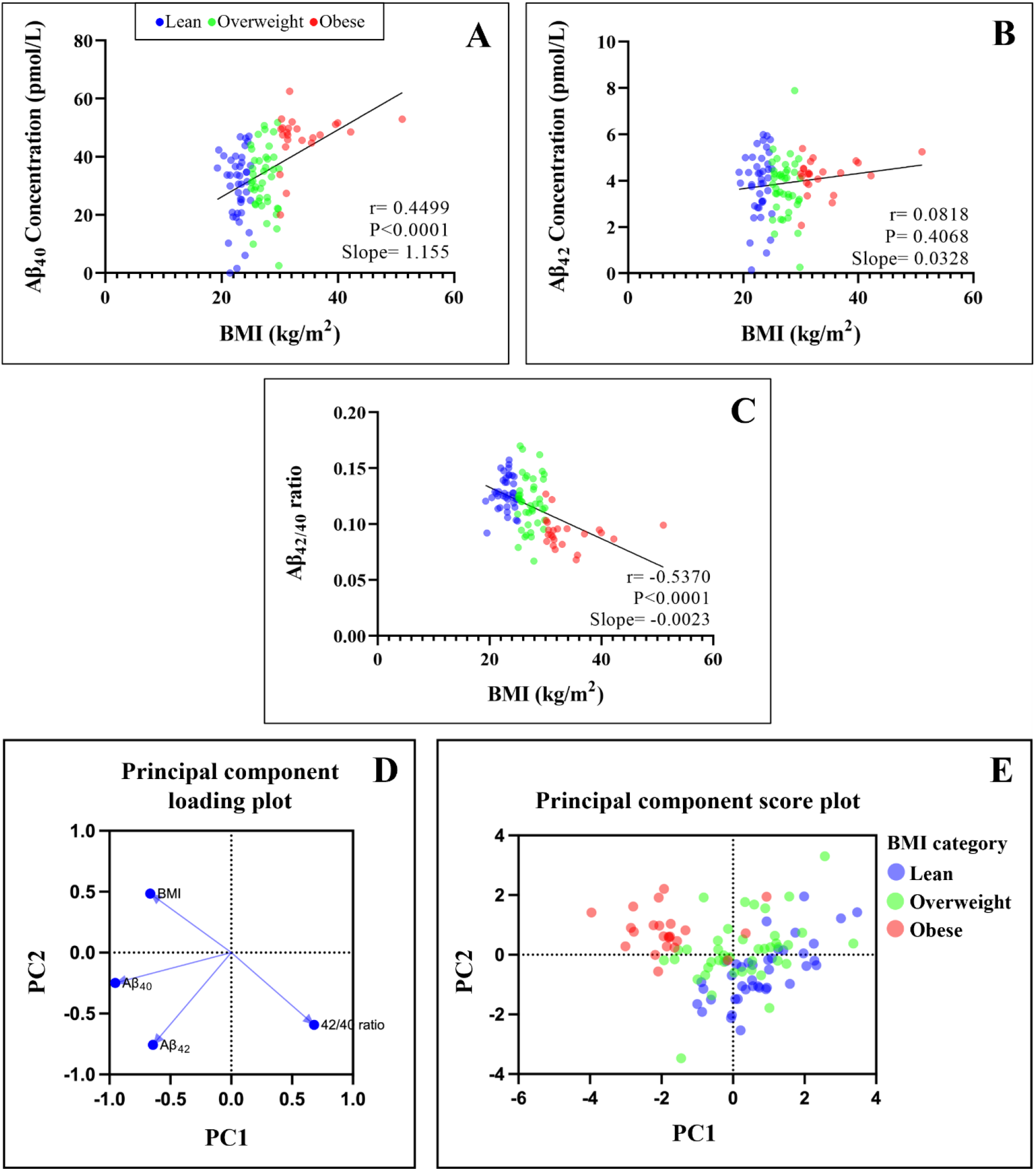
The scatter plot shows the linear correlations between BMI and plasma concentrations of A) Aβ_40_, B) Aβ_42_, and C) Aβ_42/40_ ratio. D) Shows the Principal Component Analysis loading plot of BMI, Aβ_40_, Aβ_42_ and Aβ_42/40_, and E) Principal Component Analysis score plot shows the scattering of Aβ_42/40_ in lean, overweight, and obese BMI groups.

As the average age of obese group was significantly higher compared to the lean and overweight BMI groups, and there were nearly 9-fold more men than women overall (Table 1); we used a multivariable regression model to adjust the plasma Aβ concentrations for age, age squared and sex [24]. Following adjustment, the mean plasma Aβ_40_ in the obese BMI group was ~11% greater than the lean or overweight groups; whilst plasma Aβ_40_ levels were similar between lean and overweight groups (Fig 3A). Moreover, the mean plasma Aβ_42_ concentrations were not significantly different across lean, overweight, and obese groups (Fig 3B). However, the inverse relationship between BMI and Aβ_42/40_ ratio persisted following adjustment and showed significantly lower Aβ_42/40_ in the obese group, compared to both lean and overweight BMI groups by 17.54% and 11.76%, respectively. We also note that the Aβ_42/40_ ratio in overweight trended lower than the lean group (Fig 3C).

**Fig. 3.**
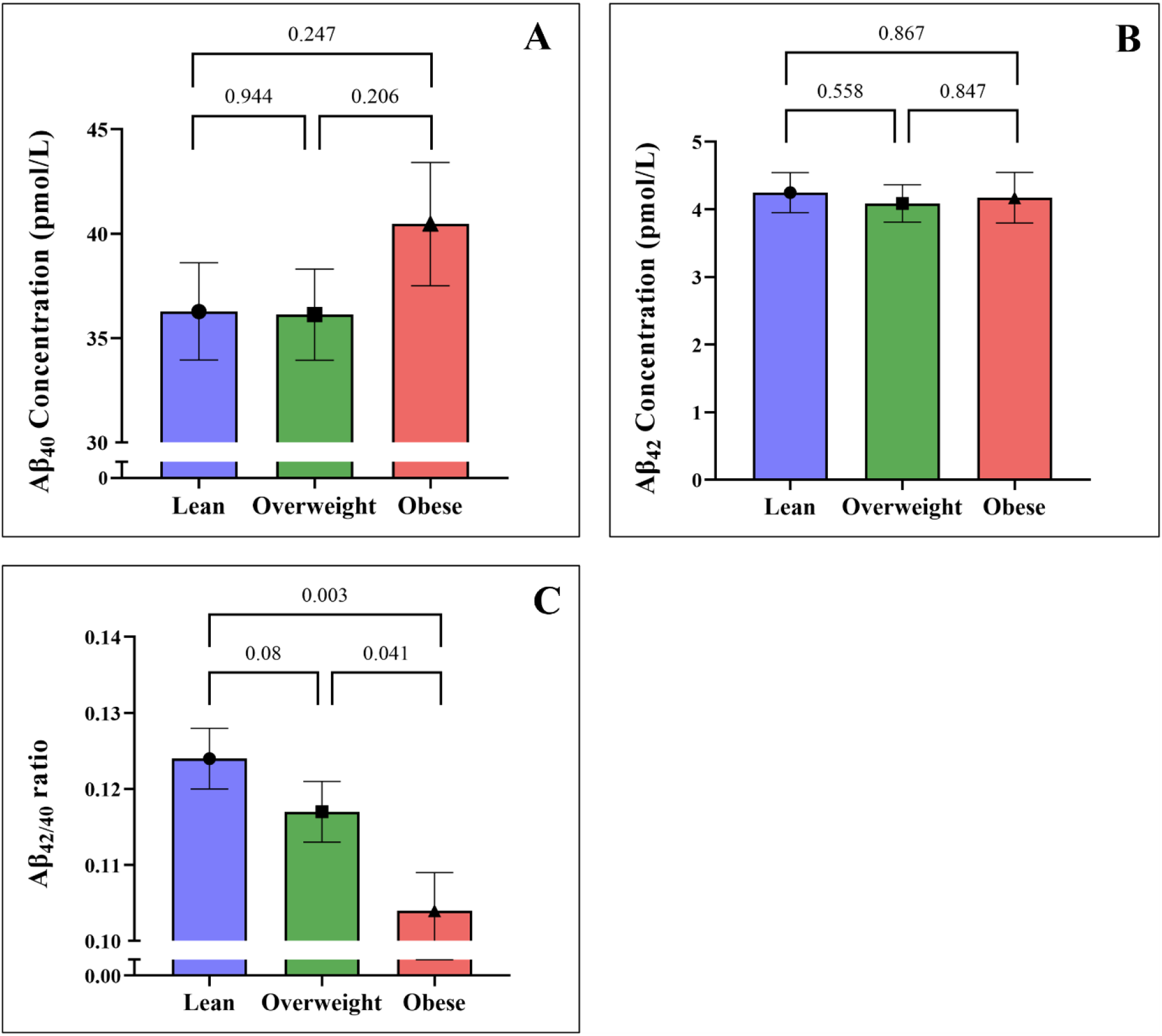
The bar graphs show adjusted plasma concentrations of A) Aβ_40_, B) Aβ_42_, and C) Aβ_42/40_ ratio in people with lean, overweight, and obese BMIs. All values are shown as mean and standard error of the mean.

To further explore the relationship between BMI and plasma Aβ homeostasis, we tested whether weight loss in participants with overweight and obese BMIs would alter plasma concentrations of Aβ.

At baseline, 2 participants had an overweight BMI, whilst the remaining 10 participants had BMIs that were classed as obese. After the 12-week weight loss intervention, participants significantly reduced their BMI by an average of 4.021% (Table 2).

**Table 2:** Age and BMI at baseline and wk-12. All values are presented as mean ± SD with range

| Variable | Baseline | wk-12 | Change (%)<sup>a</sup> | *p* value |
| --- | --- | --- | --- | --- |
| Gender (% male) | 58.33 |  |  |  |
| Age (years) | 60.17±4.22 | - | - | - |
| (range) | (52-66) |  |  |  |
| BMI (kg/m<sup>2</sup>) | 32.45 ± 3.86 | 31.18 ± 4.13 | -4.02 ± 2.25 | 0.0005 |
| (range) | (27.44-42.20) | (26.00-41.53) | (-9.58--0.43) |  |
| BMI (overweight: obese) | 2:10 | 5:7 |  |  |

<sup>a</sup>Percent change from baseline to 12-week follow-up

Plasma concentrations of Aβ_40_, Aβ_42_ and Aβ_42/40_ ratio for baseline and 12-weeks are shown in Figure 4. Following weight loss, plasma Aβ_40_ significantly decreased by 6.5% (Fig 4A). However, we note that the significant decrease in Aβ_40_ is heterogenous and predominantly driven by two participants where plasma Aβ_40_ decreased by 25.62% and 29.7%. Of these two participants, one also showed a 50.5% decrease in Aβ_42_ and a 29.6% decrease Aβ_40/42_ ratio. Whilst the other showed a 24.5% increase in Aβ_42/40_ ratio, which can be attributed to the decrease in Aβ_40_ and minimal change in Aβ_42_.

**Fig. 4.**
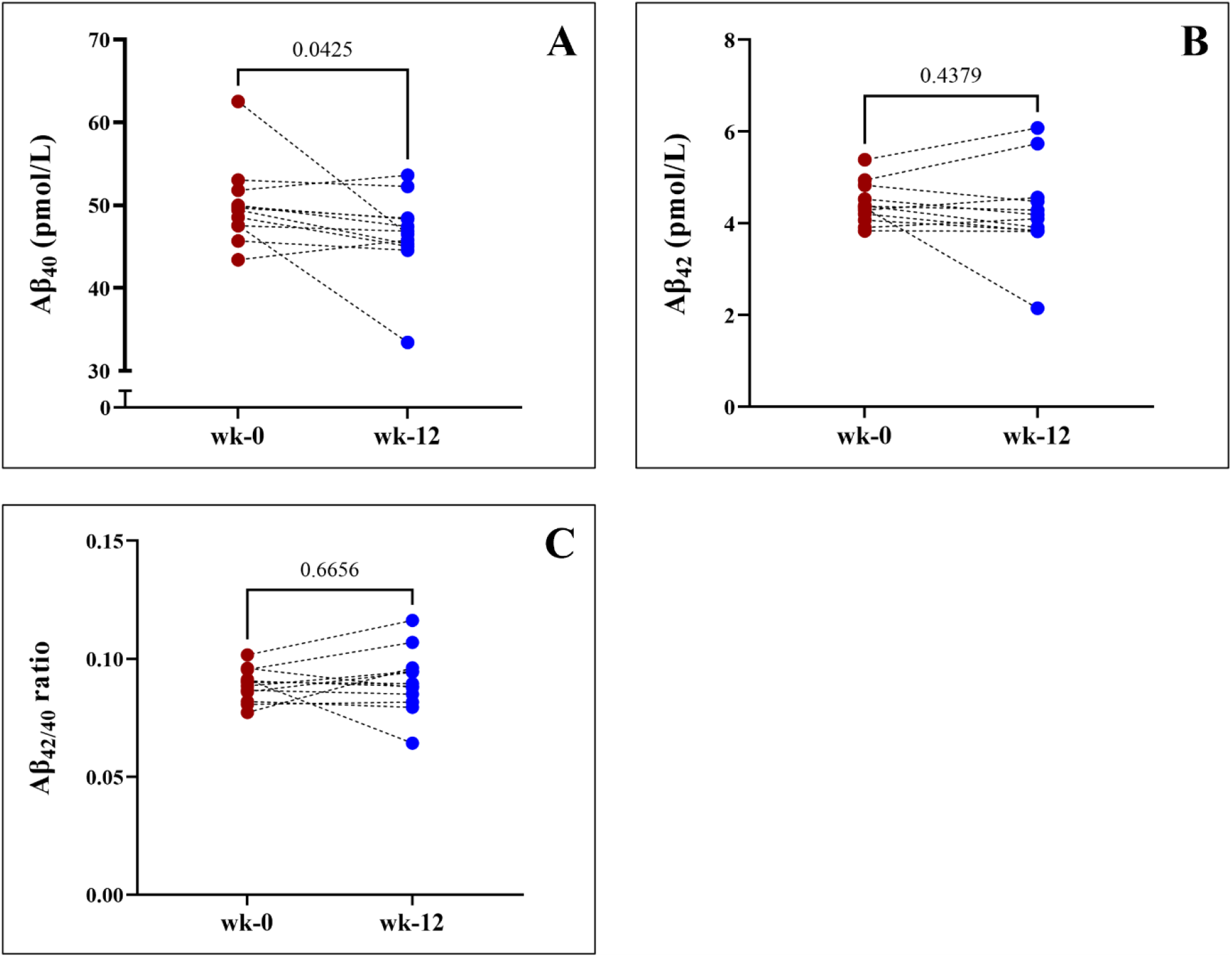
Before-after plots show the plasma concentrations of A) Aβ_40_, B) Aβ_42_, and C) Aβ_42/40_ in people who entered the calorie-restriction weight loss program.

The percent change of plasma Aβ_40_ from baseline for the remaining participants ranged from −8.35-5.45%. Plasma Aβ_42_ and the Aβ_42/40_ ratio did not show any significant changes following weight loss (Fig 4B-C).

We also performed correlation analysis between the percent change from baseline for plasma Aβ and weight lost (Fig 5). Overall, plasma Aβ_40_ shows a moderate negative correlation with percentage of weight lost, however this failed to reach statistical significance (Fig 5A). Plasma Aβ_42_ and Aβ_42/40_ ratio did not show any significant correlations with weight-loss (Fig 5B-C).

**Fig. 5.**
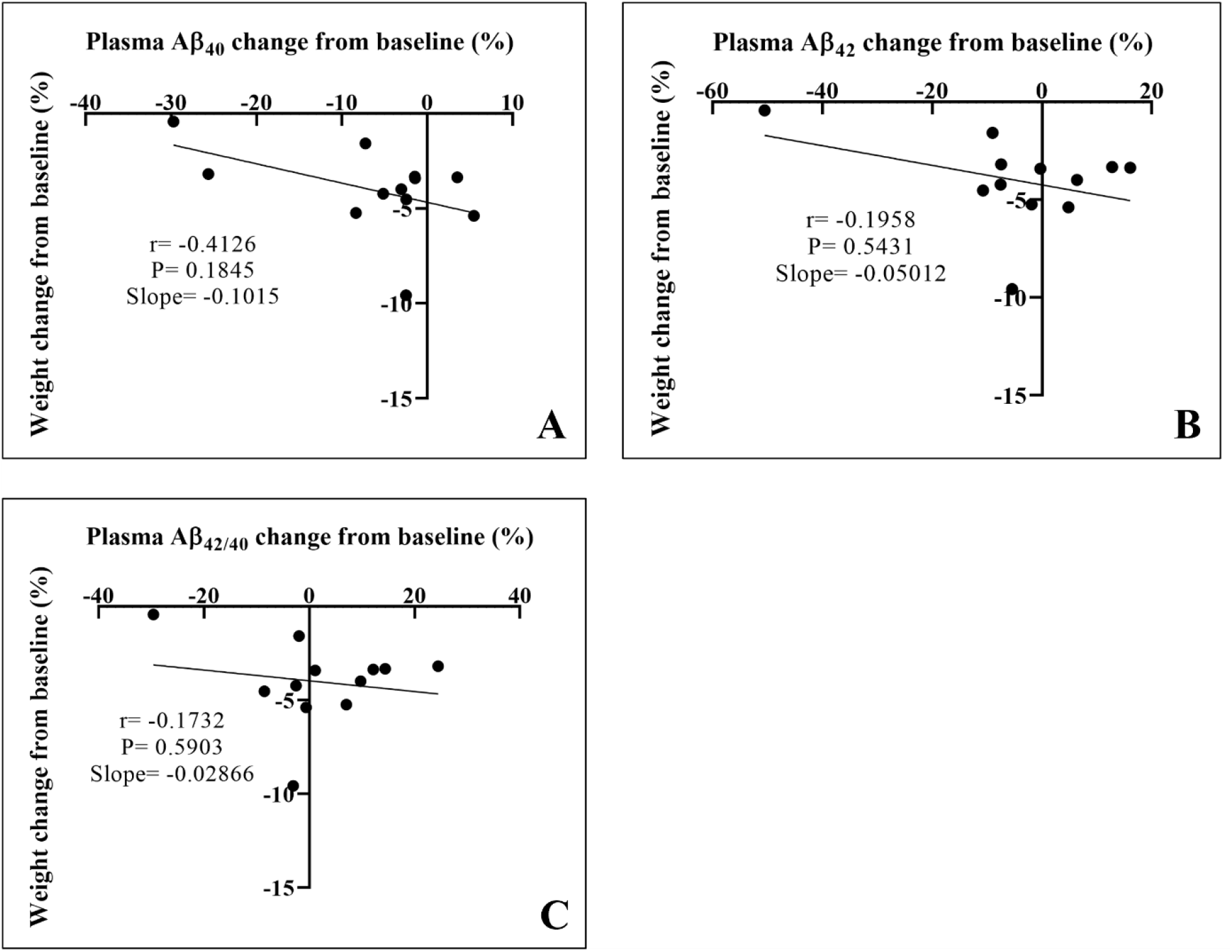
The scatter plot shows the linear correlations between the percentage of weight loss from baseline and percent change in plasma concentrations of A) Aβ_40_, B) Aβ_42_, and C) Aβ_42/40_ ratio.

## DISCUSSION

In this pilot study we investigated whether increased risk of AD in obesity is associated with plasma Aβ dyshomeostasis and examined the relationship between BMI and plasma Aβ concentrations of 106 adults.

Here, we report that plasma Aβ_40_ was significantly elevated in participants with an obese BMI compared to those in the overweight and lean groups. Consistently, plasma Aβ_40_ showed significant positive correlation with BMI, whilst plasma Aβ_42_ did not show any significant association with BMI. Nonetheless, BMI showed moderate negative correlation with Aβ_42/40_ ratio and indicated significantly lower Aβ_42/40_ ratio in the obese group compared to the people with a lean or overweight BMI. The distinct association between an obese BMI and low plasma Aβ_42/40_ ratio persisted after adjusting the data for age, age-squared and sex. Low plasma Aβ_42/40_ ratio has been consistently associated with high cerebral amyloid burden, greater cognitive deterioration, and increased risk of developing AD [12–19, 25–28]. Consequently, the low plasma Aβ_42/40_ ratio observed amongst the obese BMI group may suggest a heightened risk for AD. Collectively, our data suggests that obesity and a higher BMI may predispose people to an elevated risk of AD through disturbances in plasma Aβ homeostasis. However, the effect of obesity on plasma Aβ_42/40_ is largely understudied, and the underlying mechanisms by which obesity increases plasma Aβ_40_ and decreases in Aβ_42/40_ ratio are currently unknown.

AβPP, the precursor molecule for Aβ generation, is reported to be significantly overexpressed in the adipocytes of people with obese BMIs compared to a lean population [21, 29], indicating greater availability of Aβ biosynthesis in obesity. Additionally, preclinical findings demonstrate that in obesity, the activity of BACE1, the enzyme responsible for the extracellular cleavage of AβPP and the subsequent production of Aβ, is significantly elevated [30–32]. Moreover, pre-clinical studies report that the production and secretion into circulation of Aβ are predominantly regulated through lipid and lipoprotein metabolism, particularly cholesterol and triglyceride-rich lipoproteins [33–35]. In wild-type mice, ingestion of a diet enriched in saturated fats significantly elevated Aβ synthesis by the small intestinal enterocytes, which increased plasma Aβ concomitant with heightened non-esterified fatty acids [36, 37]. In alignment with these findings, pharmacological lipid-lowering interventions resulted in the significant reduction of plasma Aβ in wild-type mice [38–41]. These data suggest that impaired lipid metabolism and heightened plasma lipids in obesity may elevate plasma Aβ. Taken together, obesity may directly and indirectly contribute to peripheral Aβ dysregulation and lead to a low Aβ_42/40_ ratio; however, mechanistic studies are required to delineate the complete effect obesity has on Aβ homeostasis and its contribution to the AD pathological cascade.

Considering that obesity appears to disturb systemic Aβ homeostasis, we explored whether reducing BMI by calorie-restriction could influence plasma Aβ homeostasis in people with overweight and obese BMIs. Following weight loss, plasma Aβ_40_ significantly decreased, albeit heterogeneously, whilst plasma Aβ_42_ and Aβ_42/40_ ratio remained largely unaffected. Provided that our previously published findings suggest that lipid metabolism, triglyceride-rich lipoproteins and cholesterol may significantly regulate Aβ homeostasis, we speculate that reduced abundance of triglycerides and changes in cholesterol metabolism may also lead to decreased plasma Aβ_40_ [23, 36–41]. It is well documented that weight loss decreases plasma triglycerides, increases HDL cholesterol, and overall, normalises lipid profiles [23, 42, 43], and accordingly, we reported that weight loss in the present cohort decreased triglycerides and increased HDL cholesterol [23]. Interestingly, historic findings by Koudinov and colleagues indicate that apolipoproteins of HDL, namely apolipoprotein (apo) A-I, apo A-II, apo E, and apo J bind to Aβ_40_ [44]. Such findings are corroborated by recent clinical evidence whereby apo A-I appears to be a potent Aβ-binding protein [45]. Therefore, based on the current evidence, HDL may facilitate the clearance of Aβ, and we speculate that reduced Aβ_40_ following weight loss may be mediated by increased HDL.

In parallel, the adipocytes of people with obesity appear to over produce Aβ_40_, AβPP and BACE1 [21, 29–32]. Therefore, we also hypothesise that loss of adipose tissue during weight loss may reduce the availability of amyloidogenic material, and thus contribute to the reduced Aβ_40_ observed in this study. Taken together, calorie restriction induced weight loss appears to lower Aβ_40_, which may be facilitated via a HDL mediated pathway, and fat loss. Overall, the complexities of the relationship between weight loss, lipid metabolism, and Aβ_40_ is largely under studied and requires further research to determine the complete effect calorie restriction has on Aβ homeostasis in obesity.

In the present pilot study we evaluated plasma Aβ only; however, AD is a heterogenous disease with a hotly debated aetiology [46, 47], consequently numerous plasma markers of AD have been considered. Most notably, plasma phosphorylated-tau181 (p-tau18) has emerged as a robust and sensitive marker for AD. Evidence indicates that plasma p-tau181 can accurately discriminate people with AD from cognitively healthy adults, and distinguish AD from other neuropathological diseases [48]. Additionally, plasma p-tau181 can accurately predict future brain atrophy [49], cerebral deposition of amyloid, and neurodegeneration [50]. Further compelling evidence also suggests that plasma p-Tau181 may even surpass plasma Aβ_42/40_ as a predictive biomarker of AD [51, 52]. However, whether tau pathology or p-Tau181 is influenced by obesity or weight-loss is uncertain and understudied [53, 54]. In contrast, our data, along with others’ show that obesity does appear to disturb Aβ homeostasis, which may be ameliorated through weight loss, ultimately reducing risk of AD. To further substantiate these findings, we suggest that future studies employ a battery of biomarkers, including plasma p-Tau181, Aβ_42/40_ and amyloid positron emission tomography, in addition to a series of cognitive function tests.

Lastly, we acknowledge that there are demographic and methodological limitations to this pilot study. Demographically, the baseline BMI and weight loss data were obtained from limited sample sizes; the former of which was also not consistently gender balanced, although we did account for the effect of sex in our analysis. From a methodological standpoint, the weight loss intervention was conducted over a relatively short term and only modestly reduced BMI (<4.1%), which may have led to the heterogenous decrease in plasma Aβ_40_ and insignificant changes in plasma Aβ_42_ and Aβ_42/40_ ratio. Moreover, our study did not assess cognition. Considering participants from the obese BMI group were significantly older, subtle cognitive deficits and metabolic differences may have been present, which may have affected our data, although we partially addressed this matter by adjusting the data for age. Furthermore, we did not the fully account for the effect of the APOE genotype as only individuals with APOE2/2 were identified as part of the exclusion criteria. Additionally, plasma Aβ concentrations were assayed in singlets, thus any variability was not detected. We also suggest that future studies consider more sensitive methods for quantifying plasma Aβ, such as mass spectrometry-based protocols [55]. Finally, the use of BMI as an indicator of obesity has been largely criticised for not considering body composition, gender, or ethnicity [56, 57]. Therefore, in-depth studies in diverse populations which assess muscle mass and adiposity in relation to plasma Aβ would be useful to further explore how obesity influences peripheral Aβ homeostasis. Collectively, our conclusions that relate to AD development are to be interpreted circumspectly. In addition, thorough mechanistic and longitudinal studies are indeed required to substantiate the findings and interpretations of this pilot study.

In summary, our data shows that people with an obese BMI show a reduced plasma Aβ_42/40_ ratio which may translate to a heightened risk of developing AD. Importantly, we also demonstrate that lowering BMI appears to decrease plasma Aβ_40_, thus shifting it toward concentrations observed in the lean BMI group. Overall, our findings may help understand the mechanisms whereby obesity increases AD risks and offers potential preventative opportunity through weight loss.

## ACKNOWLEDGEMENTS

The authors have no acknowledgements to report.

## FUNDING

This study was funded by the National Health and Medical Research Council.

## CONFLICTS OF INTEREST

The authors have no conflict of interest to report.

## DATA AVAILABILITY

The datasets used and analysed during the current study are available from the corresponding author upon reasonable request.

## Notes

### Competing Interest Statement

The authors have declared no competing interest.

### Summary of Updates

This version of the manuscript has been revised to update grammar, flow, and clarity. Such changes were made through the entire document.

